# Jet injection potentiates naked mRNA SARS-CoV-2 vaccine in mice and non-human primates by adding physical stress to the skin

**DOI:** 10.1101/2023.02.27.530188

**Authors:** Saed Abbasi, Miki Matsui-Masai, Fumihiko Yasui, Akimasa Hayashi, Theofilus A. Tockary, Shiro Akinaga, Michinori Kohara, Kazunori Kataoka, Satoshi Uchida

**Author notes:** Correspondence to Kazunori Kataoka, Satoshi Uchida (K.K.), (S.U.). Equal contribution (Saed Abbasi, Miki Matsui-Masai).

## Abstract

Naked mRNA-based vaccines may reduce the reactogenicity associated with delivery carriers, but their effectiveness has been suboptimal against infectious diseases. Herein, we aimed to enhance their efficacy by using a pyro-drive liquid jet injector that precisely controls pressure to widely disperse mRNA solution in the skin. The jet injection boosted naked mRNA delivery efficiency in the mouse skin. Mechanistic analyses indicate that dendritic cells, upon uptake of antigen mRNA in the skin, migrate to the draining lymph nodes for antigen presentation. Additionally, the jet injector activated innate immune responses in the skin, presumably by inducing physical stress, thus serving as a physical adjuvant. From a safety perspective, our approach, utilizing naked mRNA, restricted mRNA distribution solely to the injection site, preventing systemic pro-inflammatory reactions following vaccination. Ultimately, the jet injection of naked mRNA encoding SARS-CoV-2 spike protein elicited robust humoral and cellular immunity, providing protection against SARS-CoV-2 infection in mice. Furthermore, our approach induced plasma activity of neutralizing SARS-CoV-2 in non-human primates, comparable to that observed in mice, with no detectable systemic reactogenicity.

## Introduction

The use of antigen-encoding messenger RNA (mRNA) has instigated a paradigm shift in vaccine development because of its high efficiency in vaccination, ease of mRNA sequence designing, and scalability for billions of doses per year (*1-3*). In 2020, two novel mRNA vaccines, BNT162b2 and mRNA-1273, received emergency approval for use against the coronavirus disease 2019 (COVID-19) caused by SARS-CoV-2 infection, showing >90% efficacy in preventing the symptomatic disease (*4, 5*). Both vaccines use lipid nanoparticles (LNPs) as the delivery system to protect mRNA from enzymatic degradation, facilitate its intracellular delivery, and act as an immunostimulatory adjuvant (*6, 7*). LNPs can distribute broadly to the lymph nodes (LNs), spleen, and liver following local injection in small and large mammals (*8-12*). While the distribution of LNPs to the lymphoid organs, *e*.*g*., the spleen and LNs, benefits vaccination processes, their systemic distribution raises potential safety concerns, driving extensive research to find alternative solutions (*13, 14*). Notably, reducing the incidence of reactogenic events could reduce vaccine hesitancy (*15*). In this context, enabling naked mRNA vaccines could eliminate any safety risks associated with the delivery materials and establish a new class of effective COVID-19 vaccines. However, this approach is challenging, given the multifunctional attributes of LNPs in potentiating mRNA vaccines. Indeed, the robustness of naked mRNA in protection against infectious diseases has not been well demonstrated.

Optimizing the delivery route can improve vaccination efficiency. Both BNT162b2 and mRNA-1273 are administered by intramuscular (i.m.) injections. Nevertheless, the skin represents a potent vaccination target because the epidermis and dermis have higher antigen-presenting cells (APCs) density than the muscle tissue (*16, 17*). Recent clinical trials showed that the intradermal (i.d.) delivery of BNT162b2 and mRNA-1273 achieved dose sparing at 1/10 - 1/5 of the i.m. dose, with a trend of alleviated adverse effects (*18-20*). However, using naked mRNA in i.d. vaccines remains challenging because of the poor stability and inefficient cellular uptake of mRNA in the skin (*21*). Indeed, naked mRNA i.d. vaccines yielded little or no antibody responses and only modest CD8+ cellular immunity previously (*22-24*).

The rapid delivery kinetics are crucial for internalizing naked mRNA into the cells before its degradation in the cutaneous space. To address this, herein, we employed a needle-free, pyro-drive liquid jet injector (PYRO), which facilitates the rapid internalization of macromolecules into skin cells through the instantaneous rise of liquid pressure (*25, 26*). As a result, rapid cellular internalization of mRNA following PYRO injection allowed overcoming enzymatic mRNA degradation, leading to efficient protein production from naked mRNA in the skin. Moreover, we revealed a potential function of PYRO injection as an immunostimulatory adjuvant. PYRO injection elicited localized pro-inflammatory reactions at the injection site, presumably by provoking physical stress at the injection site. Such pro-inflammatory reactions can serve as a physical adjuvant to potentiate immune responses. From a safety viewpoint, our approach of using naked mRNA restricted mRNA distribution to the injection site, avoiding systemic pro-inflammatory reactions after vaccination. Ultimately, PYRO-injected naked mRNA elicited robust vaccination effects in mice and non-human primates (NHPs) and provided a protective effect in a SARS-CoV-2 challenge test in mice.

## Results

### 1. PYRO boosts the delivery and vaccination efficiency of naked mRNA

The protein expression level in the skin following i.d. injection of *firefly luciferase* (*fLuc*)-encoding mRNA was evaluated (**Figure 1a, b**). PYRO injection induced approximately 200-fold more efficient fLuc expression at the injection site compared to manual injection with a needle and a syringe (N&S) after 4 h of injection. The fLuc expression remained high and gradually reduced in intensity over 4 days (**Figure 1a**). Despite the large difference in the protein expression efficiency, PYRO and N&S injections had similar physical appearances on the skin, showing a round, whitish bleb on the surface (**Figure 1c)**.

**Figure 1.**
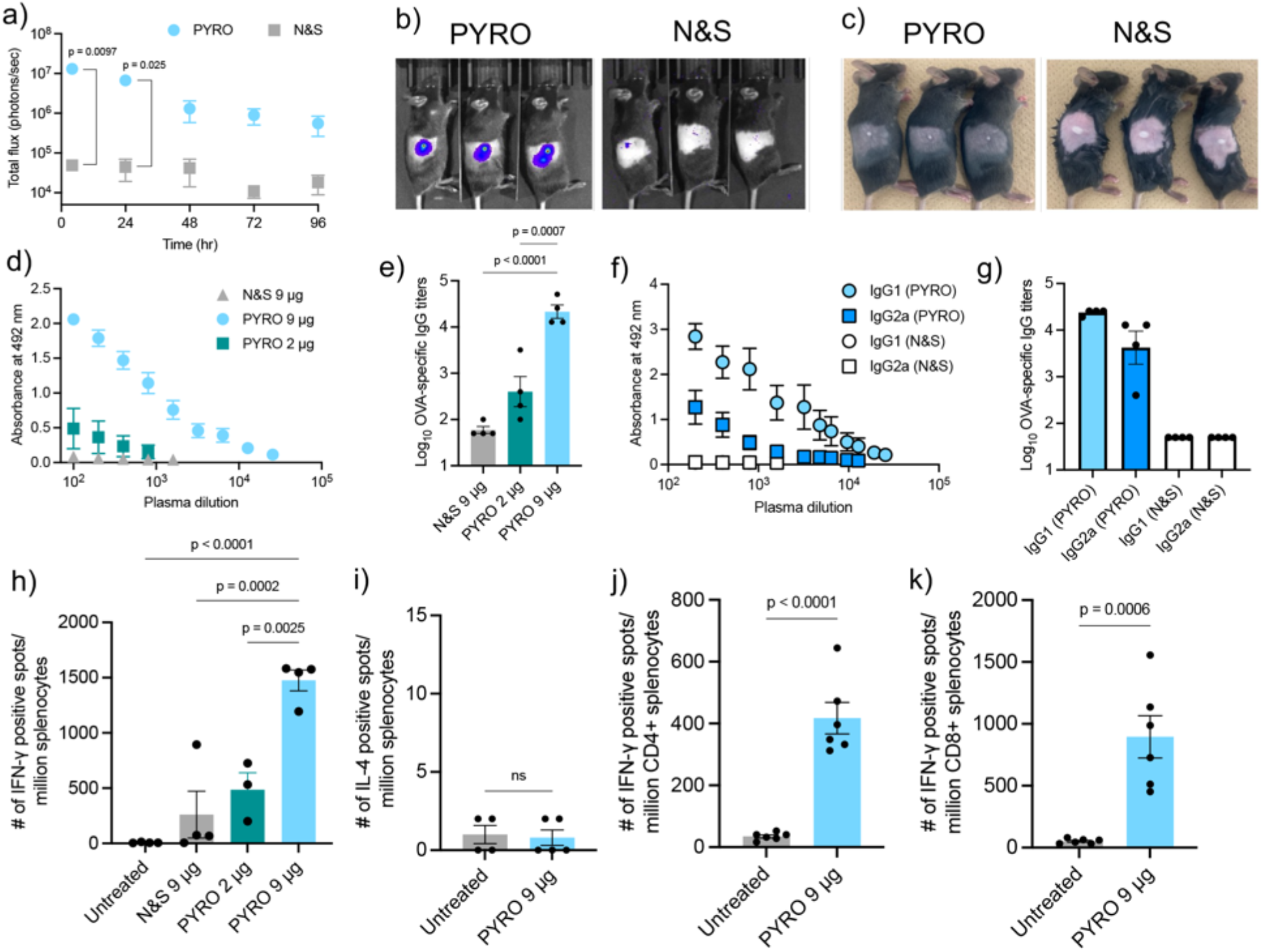
The efficiency of mRNA delivery and anti-OVA vaccination. **(a-c)** Naked mRNA encoding luciferase was i.d. injected into the right flank of mice. **(a)** Luciferase expression levels quantified by IVIS imaging. n=3. **(b)** Representative IVIS-acquired images showing luciferase expression around the injection site. **(c)** The appearance of the injection site after i.d. injection of naked mRNA using N&S and PYRO. **(d-k)** Mice were immunized with *OVA* mRNA twice at a 3-week interval, and blood and spleens were collected 2 weeks after the second dose. **(d)** OVA-specific ELISA absorbance vs. dilution curves and **(e)** Log-transformed OVA-specific IgG titers. **(f**,**g)** The isotypes of IgG produced following immunization with 9 μg *OVA* mRNA. **(f)** OVA-specific ELISA absorbance vs. dilution curves. **(g)** Log-transformed OVA-specific IgG isotype titers. n=4. **(h**,**i)** Cellular immunity as measured by INF-γ ELISpot **(h)**, and IL-4 ELISpot **(i)**. n=4. **(j**,**k)** Number of OVA-reactive INF-γ-producing CD4+ and CD8+ T cells in the spleens of vaccinated mice. n=6. Data represent the mean ± SEM. Statistical analyses were performed by one-way ANOVA followed by Tukey’s post hoc test in **(e, h)**, and unpaired Student’s t-test in **(a, i, j, k**). n.s.: nonsignificant.

We then evaluated the utility of PYRO in vaccination using naked mRNA encoding ovalbumin (OVA) as a model antigen. Mice received a prime and a boost separated by a three-week interval, each consisting of 2 or 9 μg naked *OVA* mRNA. The vaccination successfully produced OVA-specific IgG dose-dependently, with a high titer observed at 9 μg (**Figure 1d, e**). The antibody production was almost undetectable after N&S injection of 9 μg naked *OVA* mRNA. We further evaluated the isotypes of IgG produced at the 9 μg dose group. PYRO injection produced high titers of OVA-specific IgG1 and IgG2a, which were below the detection limit in the N&S group (**Figure 1f, g**). Notably, antigen-binding IgG1 and IgG2a are both needed for protection against viral infection (*27*).

The cellular immune responses of PYRO-injected naked *OVA* mRNA were also evaluated using enzyme-linked immunospot assay (ELISpot). In contrast to the absence of humoral immunity in N&S-injected mice (**Figure 1d, e**), N&S injection of naked mRNA (9 μg) induced a detectable level of OVA-specific IFN-γ-producing splenocytes in vaccinated mice, which is consistent with previous reports (*22-24, 28*). Notably, PYRO injection of naked mRNA induced a 5-fold increase in the spot number compared to N&S injection at the dose of 9 μg (**Figure 1h**). As in humoral immunity, the 9 μg dose induced higher cellular immunity than the 2 μg dose (**Figure 1h**). The splenocytes of mice injected with PYRO did not produce IL-4, a T helper 2 (Th2)-related cytokine (**Figure 1i**), but produced high levels of IFN-γ, a T helper 1 (Th1)-related molecule (**Figure 1h**). The production of both IgG1 and IgG2a and the lack of Th2 cytokines in the presence of Th1 cytokines indicate favorable humoral and cellular immune responses induced by PYRO injection of naked mRNA. Furthermore, the spleens of vaccinated mice possessed both helper CD4+ and cytotoxic CD8+ T cells reactive to OVA (**Figures 1j, k**).

### 2. PYRO-injected naked mRNA vaccine produces robust immunity and protection against the viral challenge

For SARS-CoV-2 mRNA vaccine development, mice were PYRO-injected with naked s*pike* mRNA at 2 different doses (10 and 30 μg) in a prime-boost schedule with a 3-week interval, as illustrated in **Figure 2a**. The vaccinated mice from the 2 dose groups successfully produced anti-spike-IgG one week after the boost, with the 30 μg group showing a higher IgG level than the 10 μg group (**Figure 2b**). Two weeks after the boost, mice were transduced with adenoviral vectors encoding a copy of the human angiotensin-converting enzyme 2 (ACE2) receptor by intranasal administration (*29*). Five days after the transfection, the mice were challenged with an early circulating strain of SARS-CoV-2. On day 5 after the challenge, we measured the SARS-CoV-2 viral load in the lung based on viral RNA amount evaluated by viral infectious activity (plaque forming unit, PFU) (**Figure 2c**) and quantitative PCR (qPCR) (**Figure 2d**). Compared with the untreated group, vaccination significantly reduced infectious activity in the 30 μg group (**Figure 2c**) and viral mRNA amount in the 2 dose groups (**Figure 2d**). **Figure 2e** plots the viral load (**Figure 2d**) against IgG amount **(Figure 2b**) in each mouse, revealing an inverse relationship between antibody production and viral loads in mice. This plot establishes that IgG produced by PYRO injection of naked mRNA has a protective efficacy against COVID-19 infection. In tissue sections, the lung from the unvaccinated mouse exhibited severe and diffusive inflammation and tissue damage with extensive inflammatory cell infiltration **(Figure 2f**). Intriguingly, vaccination successfully alleviated such lung inflammation in both dose groups, exhibiting only focal and modest inflammation in the lung.

**Figure 2.**
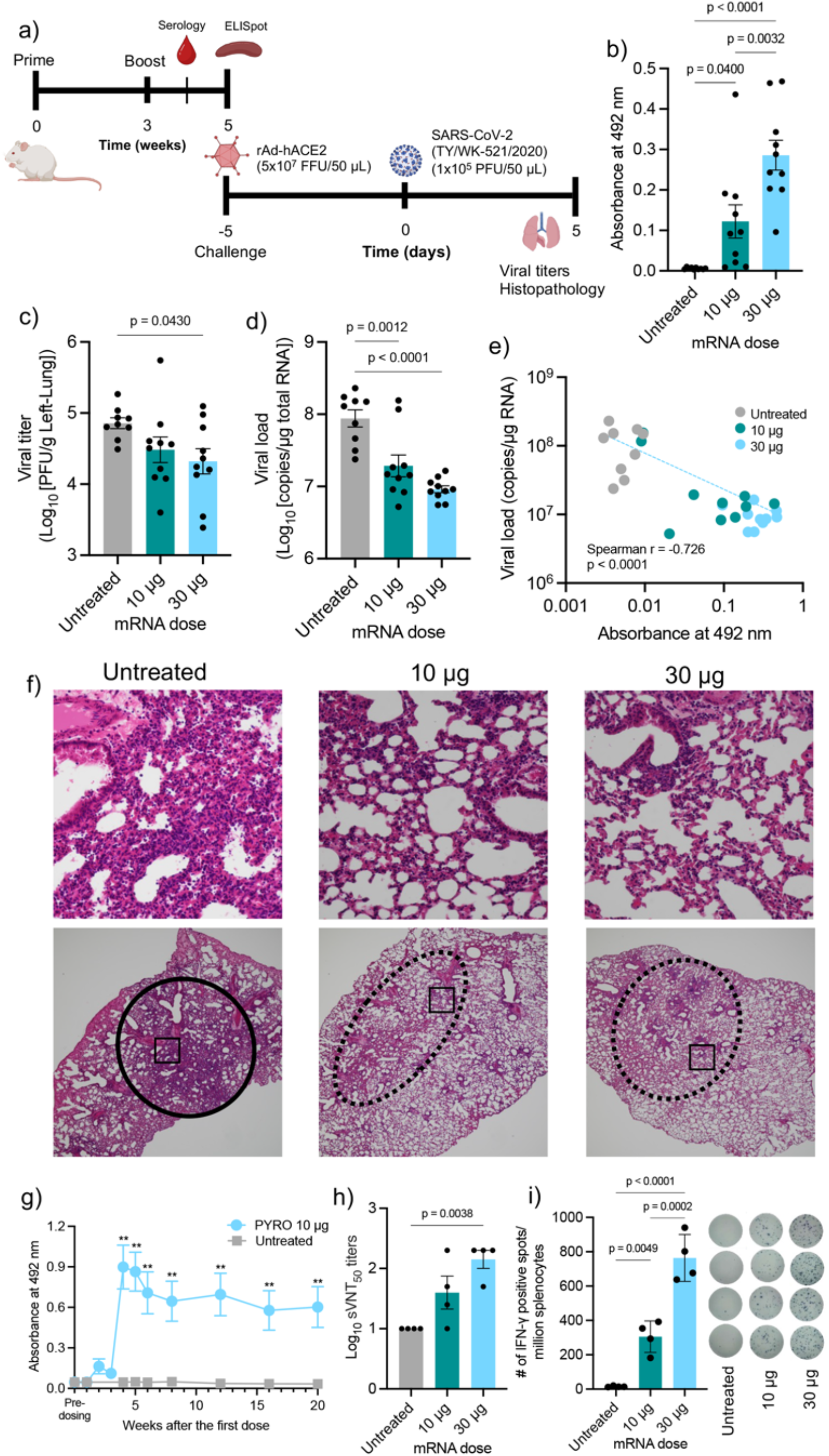
Mouse vaccination with spike mRNA and viral challenge. **(a)** Immunization and viral challenge schedule. **(b)** Spike-specific IgG ELISA absorbance at 1,000x serum dilution following PYRO injection of naked *spike* mRNA. **(c, d)** Viral challenge experiment with and without PYRO immunization with naked *spike* mRNA. Log-transformed SARS-CoV-2 viral loads in the lungs of the infected mice were measured based on viral RNA amount evaluated by viral infectious activity (PFU) **(c)** and qPCR **(d). (e)** Correlation between plasma IgG levels **(b)** and viral RNA amount in lungs **(d)**. n=10 except for the untreated group (n=9). In the untreated group, among 10 infected mice, one mouse died before the evaluation day. **(f)** H&E staining of the lung tissue sections. The upper figures show the magnification of the rectangle in the lower figures. A solid circle in the untreated group shows diffusive and severe inflammation. Dotted circles in vaccinated groups show the area with modest inflammation. **(g)** Time dependent profile of spike-specific IgG following PYRO injection of naked *spike* mRNA at the dose of 10 μg/dose. **(h)** sVNT measurement of log-transformed plasma titer of vaccinated mice required for 50% inhibition of binding between ACE2 and SARS-CoV-2 RBD. n=4. **(i)** ELISpot for spike-reactive IFN-γ splenocytes in vaccinated mice. The right image shows the appearance of the ELISpot plate. n=4. Data represent the mean ± SEM. Statistical analyses were performed by one-way ANOVA followed by Tukey’s post hoc test in all figures except **(c)**, which was followed by Dunnett’s test.

We further performed detailed characterization of vaccination effects without viral challenge. In time-dependent profiling, spike-specific IgG levels remained high 20 week after the prime, showing durable humoral immune responses after naked mRNA jet injection **(Figure 2g**). To directly demonstrate the neutralizing potential of PYRO-induced antibodies against SARS-CoV-2, mouse plasma obtained 2 weeks after the boost was subjected to a surrogate virus neutralization test (sVNT). sVNT evaluates the capability of the plasma to inhibit the interaction between the receptor-binding domain (RBD) of the SARS-CoV-2 spike protein and human angiotensin-converting enzyme 2 (hACE2) on the enzyme-linked immunosorbent assay (ELISA) plate (*30*). **Figure 2h** shows 50% inhibition titers in sVNT, revealing that PYRO-injected naked *spike* mRNA successfully inhibited the binding between spike RBD proteins and ACE2, especially at the 30 μg dose. We also evaluated the cellular immunity induced by PYRO injection of naked s*pike* mRNA. PYRO generated IFN-γ-positive splenocytes reactive with spike protein epitopes in vaccinated mice in a dose-dependent manner, with the highest spot number observed at 30 μg (**Figure 2i**). While *spike* mRNA was injected into BALB/c mice in the experiments above, C57BL/6J similarly produced spike-specific IgG titers and cellular immunity in a dose-dependent manner after PYRO injection of naked *spike* mRNA (**Supplementary Figure S1**). Notably, the induction of humoral and cellular immunity using 2 different mRNA sequences with different sizes (4,247 nt for *spike* mRNA vs. 1,437 nt for *OVA* mRNA) demonstrates the robustness of jet injection in internalizing a wide range of naked mRNA in the skin for effective vaccination.

### 3. PYRO-injected naked mRNA produces robust humoral immunity in non-human primates

Further, the utility of naked mRNA PYRO injection was tested in NHPs. Cynomolgus monkeys received PYRO injection of naked *spike* mRNA solution three times every three weeks (**Figure 3a**). The injection volume was 50 μL per dose in monkeys and 20 μL in mice. Intriguingly, PYRO injected mRNA solution distributed horizontally in NHPs, with approximately 1 cm in diameter (**Figure 3b**). In contrast, the injection in mice provided only a 2 mm-sized area of distribution (**Figure 3c**). The wide horizontal distribution of the mRNA solution in the dermal layer in monkeys may be beneficial for providing efficient contact of mRNA with APCs, because the epidermis and dermis have high APC density. Indeed, NHPs receiving 3 doses of 100 μg naked *spike* mRNA efficiently produced IgG antibodies against spike proteins (**Figure 3d)**, with high neutralization activity to inhibit the binding of RBD and ACE2 in sVNT (**Figure 3e)**. Note that sVNT titers measurement in mice and NHPs was performed using the same experimental condition, allowing us to directly compare the results obtained for these two species. Intriguingly, vaccinated NHP plasma exhibited strong neutralization potential comparable with those of vaccinated mice (**Figures 2h, 3e**). Further, NHP plasma capability to inhibit the viral infection to the cells was tested *in vitro* using an early circulating strain of SARS-CoV-2 viruses and VeroE6 constitutively expressing transmembrane protease, serine 2 (TMPRSS2) for facilitating the infection (**Figure 3f**). All three NHPs elicited efficient viral neutralization at plasma dilution of 10-fold or higher. Vaccinated NHPs showed a tendency of enhanced cellular immunity in IFN-γ ELISpot compared to the non-vaccinated control, although the difference lacks statistical significance due to the high background values in the untreated control (**Supplementary Figure S2**). As in mice, IL-4 spots were undetected (**Supplementary Figure S2**).

**Figure 3.**
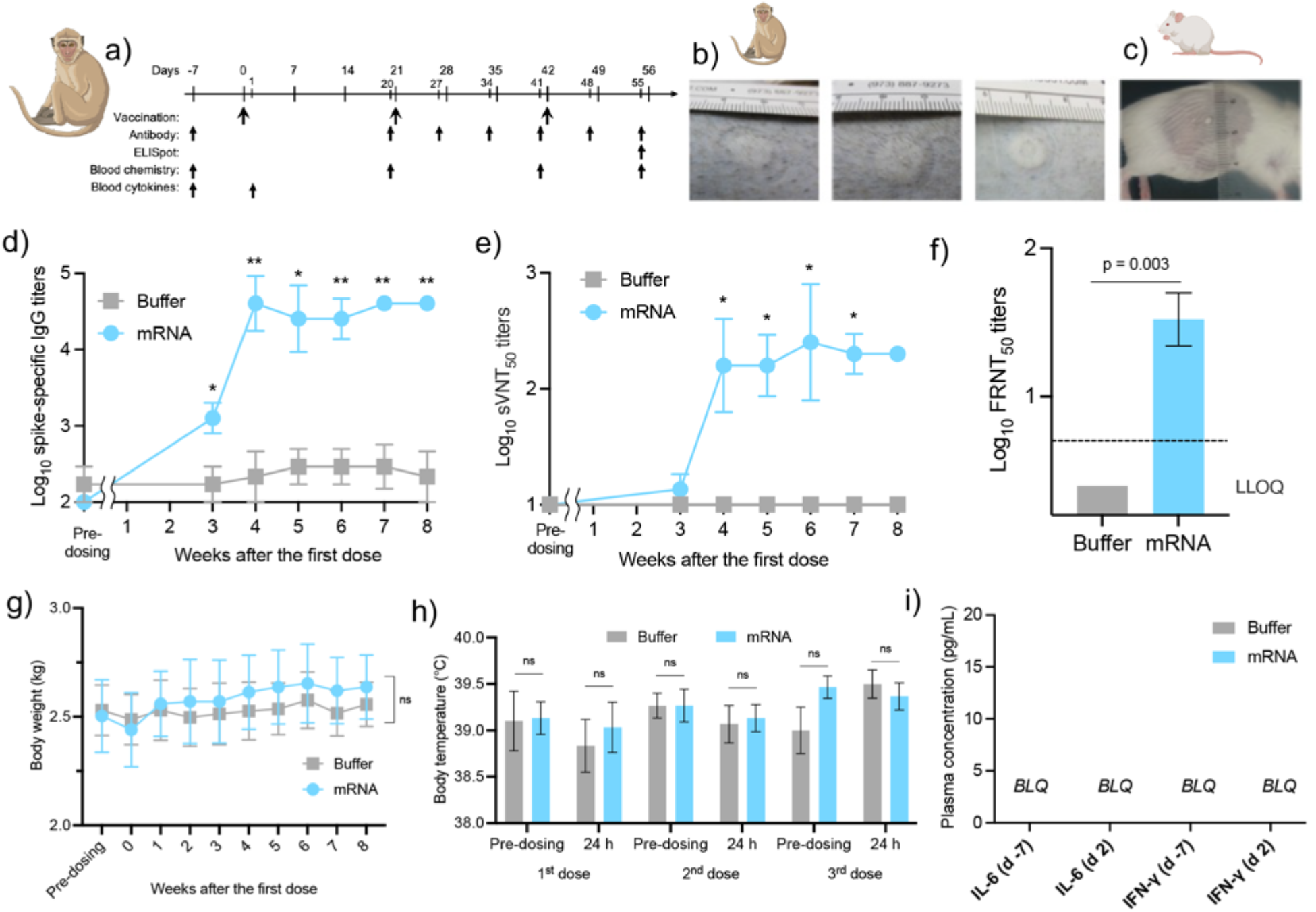
PYRO-injected naked *spike* mRNA vaccine in NHPs. **(a)** Immunization and sample collection schedule. **(b)** The appearance of PYRO injection site on the back of Cynomolgus monkeys and **(c)** mice. **(d)** Log-transformed spike-specific IgG titers in NHPs PYRO-injected with buffer or mRNA solution. **(e)** Log-transformed sVNT measurement of plasma titer required for 50% inhibition of binding between ACE2 and SARS-CoV-2 RBD. **(f)** Log-transformed plasma titer required for 50% inhibition of SARS-CoV2 infection to VeroE6/TMPRSS2 cells (FRNT_50_ titer) 8 weeks after the first dose. **(g-i)** Systemic reactogenicity in PYRO-injected NHPs receiving either buffer or *spike* mRNA. **(g)** Change in body weight. **(h)** Body temperature measured immediately before and 24 h post-dosing, after each of the 3 doses. **(i)** Plasma levels of proinflammatory cytokines before and 24 h after the first dose. LLOQ: Lower limit of quantification. n.s.: nonsignificant. BLQ: below the limit of quantification. Data represent the mean ± SEM (n=3), unpaired Student’s t-test.

NHPs were also monitored for signs of reactogenicity and toxicity throughout the 8 weeks of the study. PYRO-injected NHPs with buffer or mRNA solution showed no change in body weight **(Figure 3g)**. The body temperature 24 h after the vaccination was comparable with that immediately before vaccination in the prime or the 2 boost doses **(Figure 3h)**. The systemic release of proinflammatory cytokines such as IL-6 and IFN-γ 24 h after the prime was undetected in the blood (**Figure 3i**). Hematology and blood chemistry were comparable between mRNA-injected and buffer-injected groups (**Supplementary Tables S1, 2**). These data demonstrate high safety and robust antibody response of naked mRNA injected with PYRO injection in NHPs.

### 4. Antigen, but not mRNA, migrates to the lymph nodes

In the following sections, we performed mechanistic studies using mice to clarify why naked mRNA PYRO injection elicited robust immune responses. Locally injected vaccines need antigen trafficking to the draining LNs to generate adaptive immunity. According to previous studies, current LNP-based mRNA vaccines have 2 main modes of action in antigen presentation at the draining LNs (*1*). (i) Resident immune cells, including APCs, take up LNPs at the injection site and migrate to the draining LNs (*23*). (ii) LNPs distribute from the injection site to the draining LNs to introduce the antigen-encoding mRNA at the LNs (*12*). Here, we studied the contribution of these 2 modes of action after PYRO injection of naked mRNA. We tracked the distribution of antigens from the injection site (skin in the flank) to ipsilateral inguinal LN, a draining LN, by injecting *EGFP* mRNA as a representative of antigen-encoding mRNA. PYRO-injected naked *EGFP* mRNA induced strong EGFP expression in the immunohistochemical staining of skin sections at the injection site obtained 24 h post-injection (**Figure 4a**). The EGFP signal co-localized in part with CD11c, a DC marker, indicating mRNA expression in APCs (**Figure 4b**). Meanwhile, the remainder of the EGFP signal did not co-localize with the CD11c signal, indicating the mRNA uptake and protein expression in other cell types. In the inguinal LN, the EGFP signal was also detected 24 h post-injection (**Figure 4c**). Notably, CD11c+ DCs expressed EGFP in the LN (**Figure 4d**), suggesting successful antigen presentation at the draining LN. We also observed structural maturation of inguinal LN after vaccination using PYRO-injected naked *OVA* mRNA. H&E staining of the LN sections revealed the formation of germinal centers 2 weeks after the prime and 2 weeks after the boost (**Figure 4e, f**). This observation suggests the induction of long-lived antibody response following jet-injection of naked mRNA vaccine in the skin (*31*).

**Figure 4.**
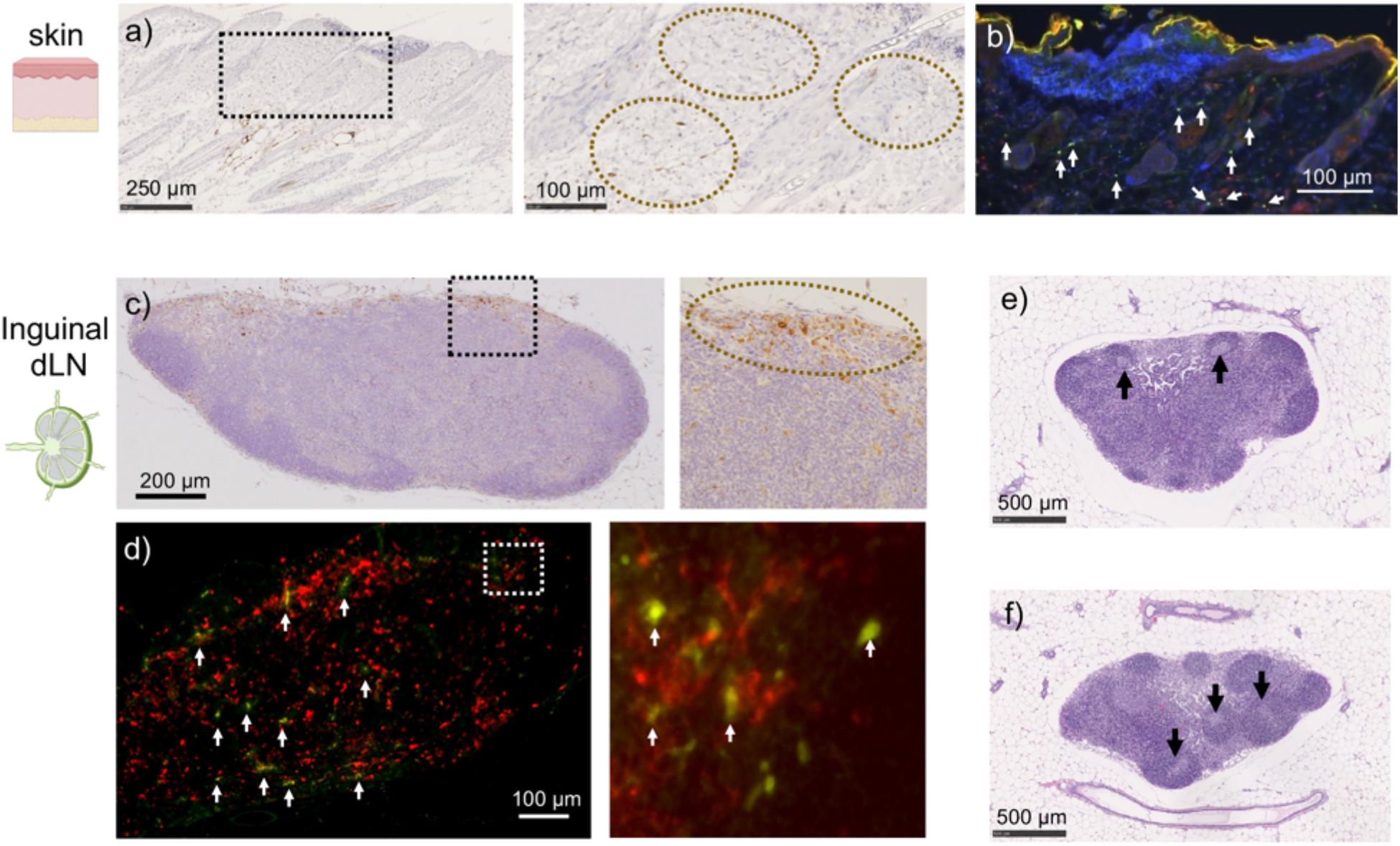
Antigen migration from skin to LNs. **(a-d**) Distribution of EGFP expression in skin (**a, b**) and ipsilateral inguinal LNs **(c, d)** 24 h after PYRO injection of naked *EGFP* mRNA into mice. **(a, c**) EGFP was stained in brown by immunohistochemistry. The right figures show the magnification of the dotted rectangle in the left figures. Dotted circles in the right figures highlight the area rich in EGFP-positive cells. **(b, d)** EGFP expression in dendritic cells was evaluated by staining EGFP in green and CD11c in red. White arrows indicate the colocalization of EGFP and CD11c. **(e, f)** H&E staining of the ipsilateral inguinal LN. Arrows indicate germinal centers. **(e)** 2 weeks. **(f)** 5 weeks.

We then tracked naked mRNA distribution to study its contribution in vaccination processes. For this purpose, the levels of injected mRNA were measured at both the inguinal and auxiliary LNs ipsilateral to the injection site by qPCR. Since naked mRNA may show rapid degradation in the extracellular and intracellular milieu (*32*), we measured its levels at early time points, 30 min and 4 h post-injection. mRNA was undetectable in both LNs 30 min and 4 h after PYRO injection of 2 and 9 μg naked *OVA* mRNA in the skin (**Figure 5a, b**). Despite the absence of intact mRNA in the LNs, DCs in the LNs became EGFP-positive (**Figure 4d**). These results suggest that DCs taking up antigen mRNA in the skin migrate to the draining LN, wherein they interact with T and B cells for antigen presentation. We next evaluated antigen mRNA distribution to other vital organs by focusing on the liver and spleen. PYRO-injected mRNA was undetectable in these organs as quantified by qPCR 4 h post-injection at 2 and 9 μg doses (**Figure 5c, d)**. This confirms that naked mRNA does not leak from its injection site after PYRO injection. Such localized mRNA may offer a safety benefit. Indeed, transcript levels of pro-inflammatory cytokines, *interleukin* (*IL)-6* and *interferon* (*IFN)-β*, after naked mRNA PYRO injection in inguinal and auxiliary LNs, liver, and spleen were comparable with those in the untreated group (**Figure 5e-l)**. This result shows that naked mRNA PYRO injection induces minimal systemic reactogenicity.

**Figure 5.**
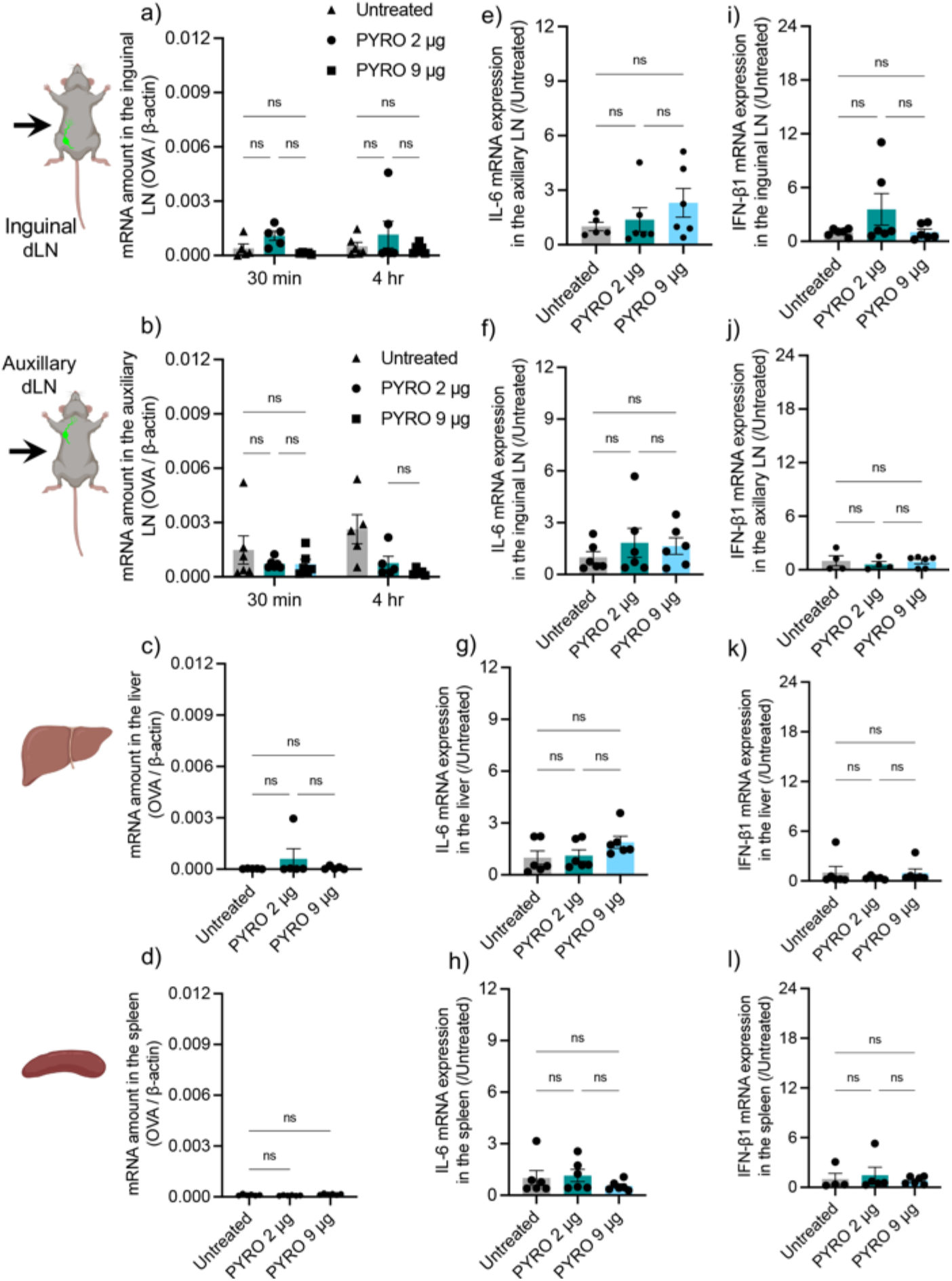
Systemic distribution of naked mRNA. **(a-d)** *OVA* mRNA levels were measured by qPCR 30 min **(a, b)** and 4 h **(a-d)** after PYRO-injection of naked *OVA* mRNA, followed by normalization to *β-actin* mRNA levels. **(e-l)** Transcript levels of *IL-6* **(e-h)** and *IFN-β* **(i-l)** were measured by qPCR 4 h after PYRO-injection of naked *OVA* mRNA, followed by normalization to untreated samples. **(a, e, i)** ipsilateral inguinal LN, **(b, f, j)** ipsilateral auxiliary LN, **(c, g, k)** liver, and **(d, h, l)** spleen. Data represent the mean ± SEM (n=5-6). Statistical analyses were performed by one-way ANOVA followed by Tukey’s post hoc test in all graphs except for **(a, b)**, which were performed using two-way ANOVA. n.s.: nonsignificant.

### 5. PYRO injections work as a physical adjuvant

Inducing high antigen expression is not sufficient to induce a robust immune response. Indeed, an mRNA delivery system that lacks an adjuvant functionality fails to induce a strong immune response even with efficient antigen expression (*33*). In the present study, naked mRNA PYRO injection exhibited a robust vaccination effect without immunostimulatory adjuvants or immunostimulatory delivery materials, such as lipid components of LNPs. This motivated us to study the immunostimulatory mechanisms after the PYRO injection of naked mRNA. We observed the immunostimulatory events at the injection site after H&E staining of the skin sections. PYRO injection of naked mRNA induced localized lymphocyte infiltration at the injection site 24 h post-injection, demonstrating its adjuvant function **(Figure 6a)**. Interestingly, similar pro-inflammatory reactions were observed after PYRO injection of the buffer without mRNA **(Figure 6b)**. These observations indicate that mRNA immunostimulatory property minimally contributes to the adjuvant effects of naked mRNA PYRO injection. Indeed, the mRNA used in this study is N1-methyl-pseudouridine-modifed and thus low immunostimulatory. In contrast to PYRO injection, N&S injection of mRNA solution or buffer failed to provide lymphocyte infiltration **(Figure 6c, d)**. These results indicate that PYRO injection, rather than mRNA properties, triggers pro-inflammatory reactions, which can function as a physical immunostimulatory adjuvant. From a safety viewpoint, it is noteworthy to mention that these pro-inflammatory reactions were localized to a few-millimeter area.

**Figure 6.**
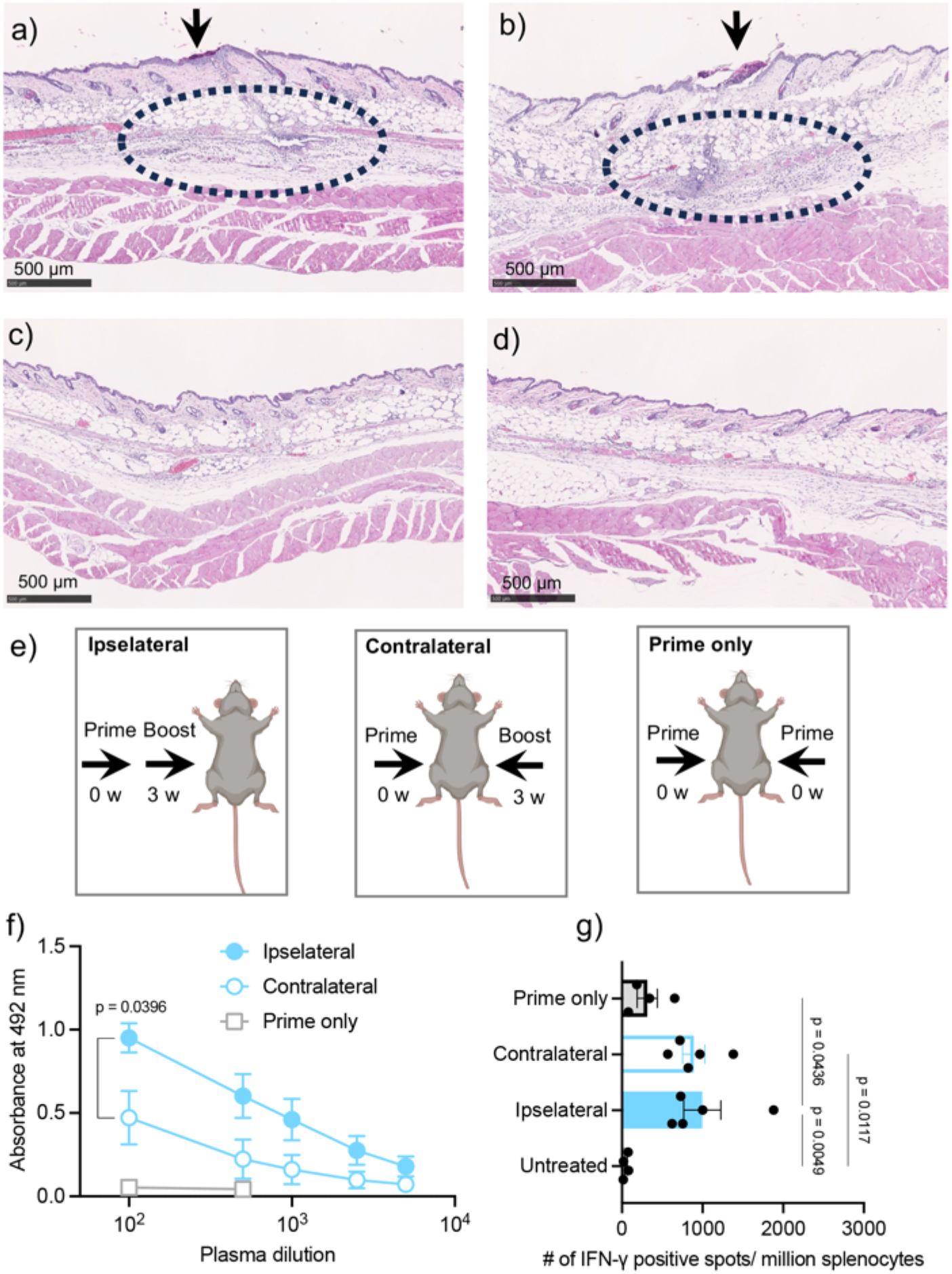
Physical adjuvant effect of PYRO. **(a-d)** H&E staining of the skin at the injection site 24 h post-injection of mRNA solution by PYRO **(a)**, buffer by PYRO **(b)**, mRNA solution by N&S **(c)**, and buffer by N&S **(d)**. Arrows represent the injection site, and the dashed circles show the infiltrations of lymphocytes. **(e)** Experimental design to study the effect of injection location on vaccine effect. **(f)** OVA-specific IgG ELISA absorbance vs. dilution curves. Data represent the mean ± SEM (n=5-6), unpaired Student’s t-test. **(g)** Cellular immunity as measured by ELISpot assay using splenocytes of vaccinated mice. Data represent the mean ± SEM (n=5-6), one-way ANOVA followed by Tukey’s post hoc test.

Considering the contribution of the local immune reactions shown in **Figure 6a**, it became reasonable to assume that the location of PYRO injections might influence the vaccination outcomes. To clarify this point, we studied the position effect of the prime and boost doses using the following two mouse groups; one group received the boost on the same side of the prime (the ipsilateral group), while the other group received the 2 doses on opposite sides (the contralateral group) (**Figure 6e**). The ipsilateral boost group induced higher amounts of OVA-specific IgG compared to the contralateral boost group (**Figure 6f**), suggesting the influence of the local immune-activated environment on the vaccination effects of the boost dose. The effect on cellular immunity was not as evident as that on humoral immunity, showing that both ipsilateral and contralateral boosting produce comparable cellular immunity levels (**Figure 6g**). We also injected mice with 18 μg naked *OVA* mRNA at a single time point (prime only) to test the effect of repetitive administration. Prime-only administration schedule failed to induce anti-OVA IgG and cellular immunity as efficiently as observed in injecting mice with 9 μg per prime and boost (**Figure 6f, g**). A previous LNP study also showed the advantage of a prime-boost protocol over a prime-only protocol in vaccination using BNT162b2 COVID-19 mRNA (*34*). In addition, despite the lack of specific recommendations on the side of injection for the prime and boost administration of COVID-19 mRNA-LNP vaccines in the clinical setting, preliminary studies were carried out by injecting the animals on the same side for both prime and boost (*35, 36*).

## Discussion

The skin is a highly immunogenic site for vaccination. Herein, we used a liquid jet injector (LJI) to facilitate mRNA delivery to the skin cells. Conventional LJI actuators, including those of the Lorenz force, are bulky and incapable of precisely controlling the injection depth (*37, 38*). In contrast, PYRO is a portable LJI operated by a miniaturized single actuator. More importantly, PYRO precisely controls the injection pressure using a bi-phasic explosion. In the bi-phasic modes, an initial explosion makes a hole in the skin across the stratum corneum, and a second explosion widely disperses the solution in the cutaneous space from the end of the hole (*25, 26*). Since optimal pressure required for wide liquid dispersion may depend on the animal species, age, and dosage, PYRO allows modulable control of injection depth and dose spreading by tuning parameters in each explosion step. Notably, we used 2 different types of explosive cartridges in this study, one is optimized for jet injection in mice and the other in NHPs. Moreover, PYRO might potentially solve several issues of N&S i.d. vaccination, including the difficulty of administering hypodermic needle injections and needle phobia (*39, 40*).

In the present study, physical forces of PYRO injection improved the vaccination effect of naked mRNA by the following two plausible mechanisms: (i) enhancing the delivery efficiency of naked mRNA (**Figure 1a**,**b**) and (ii) inducing pro-inflammatory responses at the injection site, which may serve as an immunostimulatory adjuvant (**Figure 6a-d**). Notably, PYRO provides more than 40-fold faster flow rate of injected solution than N&S injection, applying shear stress to the skin tissue (*25, 26*). The shear stress may facilitate the cellular uptake of mRNA for the former mechanism and activate danger signals in the skin to provoke pro-inflammatory responses for the latter. Regarding the latter, PYRO elicited lymphocyte infiltration into the injection site even without mRNA, whereas N&S injection of mRNA or buffer failed to induce such responses (**Figure 6a-d**). This observation indicates that physical stress, rather than mRNA immunostimulatory properties, triggers immune responses. Previous studies have also addressed related phenomena. For example, several studies reported the pro-inflammatory role of the blood-flow-derived shear stress on endothelial cells (*41, 42*), although the strength and duration of the shear stress are largely different between blood flow and PYRO-injection. PYRO instantaneously produces strong shear stress. In addition, vaccine studies of electroporation and microneedles revealed the potential roles of physical stress as a vaccine adjuvant (*43, 44*). These studies support our finding that PYRO injection plays an immunostimulatory role in vaccination. In the future study, we need to clarify the later immunization steps bridging the PYRO-driven innate immune responses to the adaptive immune responses and vaccination outcome.

From a safety viewpoint, controlling the distribution of mRNA and antigen is a critical issue of local injection. Previous studies tackled this issue by several strategies. For instance, one strategy adds miRNA target sequences to mRNA untranslated regions to prompt mRNA degradation in the off-target tissues (*45*), and another strategy develops delivery carriers with minimal systemic distribution (*13, 14*). Our unique approach of using naked mRNA utilizes mRNA nuclease susceptibility to avoid the systemic distribution of mRNA, supported by the fact that naked mRNA is rapidly degraded after systemic distribution (*46*). Indeed, qPCR measurement did not detect intact mRNA in the draining LNs, livers, or spleens following PYRO injection of naked mRNA (**Figure 5a-d**). This strategy also avoids the potential safety risks of using delivery materials, such as immunostimulatory lipids in LNPs, which distribute systemically. As a result, naked mRNA did not induce pro-inflammatory responses in the draining LNs, livers, or spleens of vaccinated mice (**Figure 5e-l**). These safety profiles of naked mRNA vaccines might solve several concerns of current mRNA vaccines, including rare cases of hepatic autoimmunity (*47, 48*) and allergic reactions (*49*). Meanwhile, systemic distribution of mRNA, especially in the lymphoid organs, *e*.*g*., the spleen and LNs, is considered to play a critical role in current LNP-based mRNA vaccines (*12*). However, naked mRNA PYRO injection provided robust immunization even without distribution to the lymphoid organs. In our mechanistic experiments using *EGFP* mRNA as a representative of antigen mRNA, EGFP-positive APCs existed in the draining LN after PYRO injection (**Figure 4d**), even without mRNA migration to the LN (**Figure 5a, b**). This result indicates that APCs that take up the antigen mRNA in the skin migrate to the draining LN, wherein APCs interact with T cells. This mechanism may contribute to efficient induction of humoral and cellular immunity by jet injection.

In a SARS-CoV-2 challenge experiment, PYRO injection of naked *spike* mRNA reduced viral load and alleviated tissue damage in mouse lungs (**Figure 2**). To the best of our knowledge, this is the first report demonstrating the protective effect of naked mRNA vaccines in an infectious disease model. Furthermore, PYRO-injected naked mRNA induced efficient humoral immunity even in NHPs with efficient ACE2 binding inhibition in sVNT comparable with that in mice **(Figures 2h, 3e**). The wide distribution of mRNA solution in the dermal layer, rich in APCs, may explain robust vaccination in NHPs (**Figure 3b**). Moreover, NHP plasma effectively prevented *in vitro* SARS-CoV-2 infection in cultured cells (**Figure 3f**). Such efficient neutralizing activity is favorable in COVID-19 vaccines, as neutralizing activity is strongly correlated with vaccine effectiveness in preventing infection of SARS-CoV-2 in the clinic (*50, 51*). Naked mRNA may have an additional advantage in storage, showing full activity when stored at temperatures as low as 4 °C in water or the lyophilized form for at least for a month (*52*). For further clinical development, we plan to study the protective effect of PYRO-injected naked mRNA against more relevant variants of SARS-CoV-2. In addition, the potency will be further optimized by tweaking other vaccine design parameters, including mRNA chemical modifications, combining different classes of immunostimulatory adjuvants, and dosing schedule.

## Materials and Methods

### 1. Materials

CleanCap® *EGFP* mRNA, N1-methyl-pseudouridine (m1Ψ)-modified *OVA* mRNA, and m1Ψ-modified *spike* mRNA with di-proline substitutions of K968 and V969 were obtained from Trilink biotechnologies (San Diego, CA, USA). The Actranza™ lab i.d. delivery device (PYRO) was purchased from Daicel Corporation (Tokyo, Japan). Ovalbumin (OVA) was purchased from Sigma Aldrich (St. Louis, MO, USA). A goat anti-mouse IgG, IgG2a, and IgG1 HRP-conjugated antibody were bought from Abcam (Cambridge, UK), R&D systems (Minneapolis, MN), and Cytiva (Tokyo, Japan). Anti-mouse IgG HRP-conjugated antibody was obtained from Cytiva in **Figure 3b**, from R&D systems in **Figure 6f**, and Abcam in the other experiments. Anti-IFNγ and anti-IL-4 ELISpot PLUS kits were purchased from Mabtech (Nacka Strand, Sweden). PepTivator Ovalbumin epitope mix was provided by Miltenyi Biotec (Nordrhein-Westfalen, Germany). PepMix™ SARS-CoV-2 (S) was obtained from JPT Peptide technologies (Berlin, Germany). Paraformaldehyde (PFA, 16%) was purchased from Alfa Aesar (Haverhill, MA, USA). SARS-CoV-2 Spike RBD-ACE2 Blocking Antibody Detection ELISA Kit was obtained from Cell Signaling Technologies.

### 2. mRNA synthesis

Luciferase DNA was constructed by cloning luciferase gene (*luc2*) (pGL4.13, Promega, Madison, WI), 120 adenine, and BsmBI cut sites into pSP73 plasmid vector. The plasmid was amplified in E. coli DH5α competent cells (Takara Bio Inc., Otsu, Japan), and then extracted and purified using a Nucleobond xtra maxi plus EF (Takara bio). Plasmids were linearized and fragmented by incubation with BsmBI overnight at 55 °C, and the desired DNA fragment was separated by gel electrophoresis and extracted using gel extraction kit (Qiagen, Hilden, Germany). The extracted DNA was further treated with T4 DNA polymerase (Takara Bio Inc., Otsu, Japan) for DNA blunting. Finally, *in vitro* transcription (IVT) synthesis of mRNA was carried out using MEGAscript™ T7 Transcription Kit (Waltham, MA, USA) with the addition of ACRA 5’ cap (Trilink biotechnologies) and m1Ψ-5’-triphosphate (Trilink biotechnologies) instead of uridine bases. The reaction was allowed to proceed for 1 h at 37 °C, and the transcribed mRNA was purified using RNeasy mini-Kit (Qiagen, Hilden, Germany). The quality of mRNA was checked using a Bioanalyzer (Agilent Technology, CA, USA).

### 3. IVIS imaging

Female C57BL/6J were obtained from Charles River Laboratories Inc., Yokohama, Japan, and used at 6-9 weeks of age. All mouse experiments are performed under the ethical guidelines of the Innovation Center of NanoMedicine (iCONM), Kawasaki Institute of Industrial Promotion (Kanagawa, Japan). A luciferase reporter assay was used to quantify protein expression following i.d. delivery of 1 μg *luc2* mRNA. For needle and syringe (N&S) injections, mouse fur was removed using Epilat cream at one flank, and 20 μL HEPES buffer (10 mM, pH 7.4) containing naked *luc2* mRNA using a 35-gauge needle (FastGene™ Nano Needle, Nippon Genetics, Tokyo, Japan). For PYRO injections, mouse fur was shaved, and 20 μL of the same buffer containing naked mRNA was injected following the manufacturer’s protocol. Protein expression was then quantified at 4, 24, 48, and 72 h post-injection using in vivo imaging system (IVIS, PerkinElmer). At each time point, imaging was performed 10 min after intraperitoneal (i.p.) injection with 200 μL of 15 mg/mL luciferin substrate (Promega). The total flux of luminescence was calculated by gating a region of interest (ROI) at the injection site.

### 4. Mouse immunization studies

*OVA* mRNA or *spike* mRNA was dissolved in HEPES buffer (10 mM, pH=7.3) and injected in the naked form. 20 μL of the solution containing a specific amount of mRNA was injected followed by a booster 3 weeks later. 2 weeks after the boost, female BALB/c or C57BL/6J mice (Charles River Laboratories Inc.) were euthanized, and blood was collected from the inferior vena cava in heparinized tubes unless specified otherwise. Blood plasma was obtained by centrifugation at 2,000 ×g for 10 min at 4 °C and stored at -80 °C until used as described in section 5. Spleens were also collected and processed for ELISpot assay as described in section 6.

### 5. Detection of mouse antibodies

Antibodies in the blood plasma were evaluated using enzyme-linked immunosorbent (ELISA). For the detection of antigen-specific antibodies, OVA or recombinant spike protein were dissolved at 2 μg/mL in carbonate buffer (50 mM, pH=9.6). 50 μL/well of protein solution was added into Clear Flat-Bottom Immuno Nonsterile 96-Well Plates (Thermo). After overnight incubation at 4 °C, the plates were washed 3 times with 0.5% w/v Tween 20 in PBS (PBS-T). A 100 μL diluent of 1% BSA and 2.5 mM EDTA in PBS-T was added to each well and further incubated for 1 h at 23 °C. After removing the diluent, 50 μL blood plasma that was serially diluted in the same diluent was added to each well. The plates were incubated overnight at 4 °C, before washing 3 times with PBS-T. A 50μL of goat anti-mouse IgG (1:8000), IgG1 (1:10000), or IgG2a (1:10000) HRP-conjugated antibodies were added to each well and incubated for an additional 2 h at 23 °C. Finally, 100 μL/well of the HRP substrate was added and incubated for 30 min at 23 °C away from light. The reaction was stopped by adding 2M sulfuric acid and measuring the absorbance of 492 nm using a plate reader (Tecan, Switzerland). Titers were determined as the highest dilution that showed absorbance optical density > 0.1.

The detection of antibodies that block the interaction between the receptor-binding domain (RBD) of the SARS-CoV-2 spike protein and angiotensin-converting enzyme 2 (ACE2) was measured by sVNT using a SARS-CoV-2 Spike RBD-ACE2 Blocking Antibody Detection ELISA Kit following the manufacturer’s protocol. Titers were determined as the highest dilution that showed binding inhibition >50%.

### 6. ELISpot assay in mice

Spleens from immunized mice were collected and disintegrated separately using a steel grid mesh with 5 mL of RPMI-1640 medium containing 10% FBS, 1 mM sodium pyruvate, 10 mM HEPES, 50 μM mercaptoethanol, and 1% penicillin/streptomycin. The suspension was then passed through a 40 μm nylon mesh (Cell strainer, Falcon) to form a single-cell suspension. Splenocytes were seeded at a density of 2.5× 10^5^ cell/well in anti-IFNγ or anti-IL-4-ELISpot 96 well plates and stimulated by the addition of 10 μL epitope mixture dissolved in PBS (0.025 μg/well OVA epitope mix and 0.2 μg/well spike epitope mix). The plates were incubated at 37 °C and 5% CO_2_ overnight. The next day, the plates were washed and treated according to the manufacturer’s protocol. Spots were counted on an ELISpot plate reader (AID GmBH, Germany).

Manual MACS® Magnetic Separator (Miltenyi Biotec) was used to separate CD4+ and CD8+ T lymphocytes from the cell suspension following the manufacturer’s instructions. CD4+ or CD8+ splenocytes were seeded at a density of 2.5 × 10^5^ cell/well in anti-IFNγ ELISpot 96 well plates. To detect the number of OVA-reactive CD4+ or CD8+ cells, OVA epitopes were presented on the surface primary DC obtained from donor mice. Primary DCs were harvested from mice femur and cultured according to a previous protocol (*53*). Primary DCs were cultured in 12-well plates at a density of 2.5 × 10^5^ cell/well and incubated with 1 μg OVA epitope mix overnight. 50 μL containing 15,625 primary DCs pulsed with OVA epitope were then added to CD4+ or CD8+ cells splenocytes and further incubated overnight. Plates were further treated and spots were counted as described above.

### 7. Viral challenge experiment

The SARS-CoV-2 TY/WK-521/2020 strain (GISAID ID: EPI_ISL_408667) was used. This strain was provided by Dr. Masayuki Saijo, Mutsuyo Takayama-Ito, and Masaaki Satoh (Department of Virology 1, National Institute of Infectious Diseases), and was subcultured in Vero E6/TMPRSS2 cells (JCRB1819, Japanese Collection of Research Bioresources (JCRB) Cell Bank, National Institute of Biomedical Innovation, Health and Nutrition, Osaka, Japan) and grown in DMEM (Nissui Pharmaceutical Co. Ltd., Tokyo, Japan) supplemented with 10% inactivated fetal bovine serum, penicillin (100 units/mL), streptomycin (100 μg/mL), and G-418 (1 mg/mL). All procedures using SARS-CoV-2 were performed in biosafety level 3 facilities by personnel wearing powered air-purifying respirators (Shigematsu Co., Ltd., Tokyo, Japan).

Female Balb/c mice (Japan SLC inc., Shizuoka, Japan) were vaccinated twice every three weeks. Two weeks after the boost, mice were infected with 1 × 10^5^ PFU/50 μL/animal of SARS-CoV-2 early circulating strain (TY/WK-521). Five days before virus infection, mice were inoculated intranasally with 5 × 10^7^ FFU/animal of rAd5 hACE2 (*29, 54*). The body weight of each mouse was monitored, and the loss of 30% of initial body weight was defined as the endpoint for euthanasia. All animals were euthanized at 5 days post infection, and blood and lungs were then collected.

The lungs of mice were homogenized using a Multi-beads shocker (Yasui Kikai, Osaka, Japan) in 9 volumes of Hanks’ Balanced Salt Solution (Gibco #14025-092). Total RNA was extracted from lung homogenates using an RNeasy Mini kit (Qiagen) according to the manufacturer’s instructions. The levels of RNA corresponding to the N protein-encoding gene of SARS-CoV-2 were measured using the TaqMan Fast Virus 1-step Master Mix (Thermo Scientific). Each 20-μL reaction mixture contained 5.0 μL of 4× TaqMan Fast Virus 1-Step Master Mix, 0.25 μL of 10 μM probe, 1.0 μL each of 10 μM forward and reverse primers, 7.75 μL of nuclease-free water, and 5.0 μL of nucleic acid extract. Amplification was carried out in 96-well plates using a CFX-96 cycler equipped with CFX Maestro software (Bio-Rad, Hercules, CA, USA). The thermocycling conditions were as follows: 5 min at 50 °C for reverse transcription, 20 s at 95 °C for the inactivation of reverse transcriptase and initial denaturation, and 45 cycles of 5 s at 95 °C and 30 s at 60 °C for amplification. Each run included a no-template control reaction as well as reactions intended to provide a standard curve. The latter used *in vitro* transcribed RNA of the N protein-encoding gene (at 10^0^, 10^1^, 10^2^, 10^3^, 10^4^, 10^6^, and 10^8^ copies/reaction); this template was generated from the cDNA of SARS-CoV-2 AI/I-004/2020 using the T7 RiboMAX Express Large Scale RNA Production System (Promega, Madison, WI, USA). The primers and probe used to detect the WK-521 strain were as follows: forward primer, 5’-GACCCCAAAATCAGCGAAAT-3’; reverse primer, 5’-TCTGGTTACTGCCAGTTGAATCTG-3’; and probe, 5’-(FAM)-ACCCCGCATTACGTTTGGTGGACC-(BHQ-1)-3’.

The infectious SARS-CoV-2 titer was determined using a standard plaque assay. Briefly, the left lung lobe from each mouse was weighed and homogenized in nine volumes of Hanks-balanced saline solution (Gibco #14025-092) using a Multi-Beads Shocker. The homogenates were centrifuged at 3000 × g for 10 min at 4ºC. The supernatant was collected and stored at −80ºC until use. Serial 10-fold dilutions of the supernatant or virus solutions (100 μL per each well) were inoculated onto confluent monolayers of Vero E6/TMPRSS2 cells in 6-well plate and incubated at 37 °C for 1 h. Unbound viruses were removed by washing the cells with DMEM. The cells were then overlaid with DMEM containing 10% inactivated fetal bovine serum and 0.6% agarose (Sigma-Aldrich, St. Louis, MO, USA). After 48 h of incubation at 37°C, the cells were fixed in 10% neutral buffered formalin and stained with 1% crystal violet. The titer of SARS-CoV-2 was defined as plaque-forming units per gram of lung tissue (PFU/g lung) or PFU per milliliter (PFU/mL). The detection limit is 100 PFU/g lung for the lung homogenates or 10 PFU/mL for virus stock solutions.

In Lung histopathology analyses, mouse lung samples were fixed in 10% neutral buffered formalin, embedded in paraffin, and cut into 4-μm-thick sections, which were mounted on glass slides. Tissue sections were stained with hematoxylin and eosin and observed by light microscopy.

### 8. NHP immunization studies

NHP experiments were performed by Ina Research Inc. (Nagano, Japan) under Act on Welfare and Management of Animals and animal experimental guidelines (Nagano, Japan) after being approved by Institutional Animal Care and Use Committee in Ina Research Inc. Cynomolgus monkeys (female, 3 – 4 years old) received 50 μL of lactated Ringer’s buffer or mRNA solution (100 μg/mouse) at a single position at their back in each prime and boost dosing. ELISA assay was performed as described in section 7 except for using Goat Anti-Monkey IgG H&L (HRP) for detecting monkey IgG. Titers were determined as the highest dilution that showed absorbance optical density > 0.15. sVNT was performed as described in section 5. Neutralization assay of NHP plasma against SARS-CoV2 was performed by Bio Medical Laboratories. Inc. Briefly, the diluted plasma was incubated with an early strain of SARS-CoV2 at the final viral concentration of 1 TCID_50_ (median tissue culture infectious dose) / μL for 1 h at 37 °C. Then, the solution was added to VeroE6 cells constitutively expressing transmembrane protease, serine 2 (TMPRSS2) for 5-day incubation. Finally, the cell was stained with crystal violet for evaluating the cytopathic effect of SARS-CoV2. Peripheral blood mononuclear cells (PBMCs) were collected for ELISpot assays using ELISpot Plus: Monkey IFN-γ (HRP) and ELISpot Plus: Human IL-4 (HRP) kits (Mabtech). 2 × 10^5^ PBMCs seeded onto 96 well plates in the kits were incubated with 1 μg of PepMixTM SARS-CoV-2 (Spike Glycoprotein) for 20 h followed by counting spot numbers using IMMUNOSPOT S6 Versa (Cellular Technology Ltd., Cleveland, OH). Safety analyses were performed using XN-2000 (Sysmex, Kobe, Japan), CA-510 (Sysmex) and flow cytometer (FACS Canto II, Becton, Dickinson and Company, Franklin Lakes, NJ) for measuring blood cell counts and coagulation functions, type 7180 auto analyzing machine (Hitachi, Tokyo, Japan) for blood chemistry analyses, and BDTM Cytometric Beads Array (CBA) Non-Human Primate Th1/Th2 Cytokine Kit (Becton, Dickinson and Company) for blood cytokine measurement.

### 9. Histopathology and microscopy

For the observation of protein expression in the skin following injection of 9 μg *EGFP* mRNA, around 1 cm^2^ of the skin surrounding the injection site was excised and fixed in 4% PFA in PBS for 24 h. The skin was then cut into several strips each around 2 mm in width, immersed in paraffin, and cooled down to form blocks for sectioning. Draining lymph nodes proximal to the injection site were also collected 24 h post-injection and treated the same as described above. The samples were sliced at 4 μm thickness, stained with hematoxylin and eosin (H&E), and observed under the optical microscope. EnVision system (DAKO) was applied for immunofluorescent staining of the paraffin-embedded samples with 5 min autoclaving at 121 °C under pH 9.0. Overnight incubation was performed for labeling dendritic cells (DCs) by a rabbit anti-CD11c antibody (Cell Signaling, #97585, 500× dilution) and EGFP by goat anti-GFP antibody (Abcam, ab5450, 1000× dilution). On the next day, the sections were washed and a mixture of secondary antibodies composed of Alexa Fluor 568 rabbit anti–goat IgG and Alexa Fluor 488 goat anti–rabbit IgG (Invitrogen) were applied to labeled CD11c and EGFP, respectively. The sections were incubated for 1 h at room temperature, washed, and observed under the fluorescent microscope (Keyence, Osaka, Japan) after the addition of DAPI to stain the cell nuclei. For the observation of lymph node maturation and pro-inflammatory reactions in the skin, 10 μg of spike mRNA was injected and H&E staining was performed as described above.

### 10. Quantitative PCR

For the quantification of *OVA* mRNA delivered to mouse organs following injection in the skin, lymph nodes, the liver, and spleen were extracted and homogenized by a Multi-beads shocker at 2,000 rpm for 30 sec. Total RNA was extracted from the homogenates using an RNeasy mini kit (Qiagen, Hilden, Germany). After the removal of genomic DNA by enzymatic degradation, complementary DNA (cDNA) was obtained by reverse transcription using a ReverTraAce with a gDNA remover kit (TOYOBO, Osaka, Japan). The products were run on a 7500 fast real-time PCR (Applied Biosystems) using FastStart Universal SYBR Green Master kit. For the quantification of *OVA* mRNA, a forward primer: GAACCAGATCACCAAGCCCA, and a reverse primer: GTACAGCTCCTTCACGCACT were used. Proinflammatory cytokines in these organs were also measured by quantitative PCR (qPCR), as described above, using the following primers. IL-6 (Mm00446190_m1) : 4331182, IFN (Mm00439552_s1) : 4331182, and B-Actin:4352933E. Data were analyzed with a 2^−ΔCt^ method using the β-actin gene as an endogenous control housekeeping gene. The pro-inflammatory cytokines data was presented after normalization to untreated samples.

### 11 Data representation

The statistical significance between the two groups was analyzed using an unpaired, two-tailed Student’s *t*-test. Multiple comparisons between three or more groups were performed using one-way ANOVA followed by Tukey’s post hoc test. Comparison against NT samples was performed using Dunette’s test. A statistically significant difference was set at *p* < 0.05.

## Acknowledgments

This work was supported by the Center of Innovation Program (COI) (K.K.) and COI accelerating support (S.U.) from Japan Science and Technology Agency (JST), Grants-in-Aid for Challenging Research (Pioneering) [18H05378 to K.K.], and for Scientific Research (A) [21H04962 to S.U., 21H04967 to K.K.] from the Ministry of Education, Culture, Sports, Science and Technology, Japan (MEXT), Strategic Center of Biomedical Advanced Vaccine Research and Development for Preparedness and Response (SCARDA) [233fa827XXXh0002 to S.U.], Leading Advanced Projects for Medical Innovation [21gm0010008s0101 to S.U.], Research Program on Emerging and Re-emerging Infectious Diseases [21fk0108620h0001 to S.U.] from Japan Agency for Medical Research and Development (AMED) and the Program for Promotion of Fundamental Studies in COVID-19 of the Tokyo Metropolitan Government. We would like to thank Yuki Sato, Keisuke Nagao, Yuki Tada (iCONM), Satomi Nakagahara, Yumi Fujii (NANO MRNA Co., Ltd.), Ayumi Sumiishi, Namiko Kondo and Kaoruko Kojima (Kyorin University School of Medicine) for their technical assistance, Dr. Hiroaki Shimoyamada (Kyorin University School of Medicine) for his consultation about histological evaluation, Dr. Katsumi Takaba (Skypatho) for his consultation about NHP experiments, Daicel Corporation for the assistance in jet injection and Ina Research Inc. for performing NHP experiments, Drs. Masayuki Saijo, Mutsuyo Takayama-Ito, and Masaaki Satoh of the Department of Virology 1 and NIID for sharing the SARS-CoV-2 virus.

