## Supplementary Information for "Jet injection potentiates naked mRNA SARS-CoV-2 vaccine in mice and non-human primates by adding physical stress to the skin"

Supplementary Table S1. Hematology markers following repeated intradermal injections in Cynomolgus Monkeys using PYRO

| | Injection | Day -7 | Day 20 | Day 41 | Day 55 |
|---|---|---|---|---|---|
| RBC ($10^6$/μL) | Buffer | 5.23 ± 0.23 | 5.16 ± 0.09 | 5.02 ± 0.19 | 5.04 ± 0.06 |
| | mRNA | 5.51 ± 0.22 | 5.34 ± 0.11 | 5.08 ± 0.19 | 5.06 ± 0.09 |
| HGB (g/dL) | Buffer | 13.07 ± 0.41 | 12.80 ± 0.10 | 12.43 ± 0.35 | 12.43 ± 0.15 |
| | mRNA | 14.13 ± 0.44 | 13.60 ± 0.17 | 13.03 ± 0.42 | 13.00 ± 0.21 |
| HCT (%) | Buffer | 42.87 ± 1.60 | 42.27 ± .72 | 41.67 ± 1.68 | 40.93 ± 0.61 |
| | mRNA | 45.03 ± 1.57 | 43.67 ± 0.64 | 42.47 ± 1.69 | 42.20 ± 0.74 |
| MCV (fL) | Buffer | 25.03 ± 0.41 | 24.80 ± 0.38 | 24.80 ± 0.40 | 24.63 ± 0.38 |
| | mRNA | 25.70 ± 0.67 | 25.50 ± 0.64 | 25.73 ± 0.65 | 25.73 ± 0.82 |
| MCH (pg) | Buffer | 82.00 ± 0.53 | 81.87 ± .09 | 83.03 ± 0.70 | 81.13 ± 0.37 |
| | mRNA | 82.00 ± 1.80 | 81.87 ± 1.05 | 83.67 ± 0.91 | 83.47 ± 2.12 |
| MCHC (g/dL) | Buffer | 30.50 ± 0.29 | 30.30 ± 0.44 | 29.87 ± 0.38 | 30.40 ± 0.47 |
| | mRNA | 31.40 ± 0.19 | 31.17 ± 0.44 | 30.70 ± 0.53 | 30.83 ± 0.37 |
| RET (%) | Buffer | 1.19 ± 0.20 | 1.19 ± 0.19 | 1.73 ± .44 | 1.42 ± 0.26 |
| | mRNA | 1.28 ± 0.21 | 1.86 ± 0.36 | 2.29 ± 0.36 | 1.67 ± 0.29 |
| RET ($10^9$/L) | Buffer | 61.17 ± 8.15 | 61.10 ± 8.98 | 85.40 ± 19.87 | 71.33 ± 12.55 |
| | mRNA | 70.33 ± 8.71 | 98.17 ± 16.98 | 116.17 ± 15.50 | 84.33 ± 13.52 |
| PLT ($10^3$/μL) | Buffer | 400.0 ± 54.0 | 376.3 ± 62.4 | 395.33 ± 63.3 | 391.7 ± 46.3 |
| | mRNA | 321.7 ± 49.3 | 292.0 ± 58.6 | 313.3 ± 62.2 | 324.0 ± 46.3 |
| WBC ($10^3$/μL) | Buffer | 11.47 ± 0.37 | 13.52 ± 1.54 | 10.92 ± 0.79 | 9.69 ± 0.27 |
| | mRNA | 8.06 ± 0.67 | 11.77 ± 1.22 | 10.08 ± 0.75 | 9.97 ± 1.07 |
| NEUT (%) | Buffer | 28.07 ± 5.17 | 39.53 ± 7.34 | 35.53 ± 1.49 | 36.63 ± 4.78 |
| | mRNA | 34.17 ± 2.68 | 47.90 ± 8.42 | 40.00 ± 1.72 | 47.57 ± 8.66 |
| LYMPH (%) | Buffer | 68.37 ± 4.83 | 56.50 ± 6.54 | 60.60 ± 1.14 | 59.83 ± 3.82 |
| | mRNA | 61.13 ± 3.20 | 48.33 ± 7.77 | 55.00 ± 1.58 | 48.80 ± 8.12 |
| MONO (%) | Buffer | 2.87 ± 0.33 | 3.30 ± 0.87 | 3.33 ± 0.44 | 2.97 ± 0.87 |
| | mRNA | 3.97 ± 0.64 | 3.33 ± 0.85 | 4.30 ± 0.50 | 3.13 ± 0.81 |
| EO (%) | Buffer | 0.57 ± 0.09 | 0.53 ± 0.15 | 0.40 ± 0.10 | 0.47 ± 0.15 |
| | mRNA | 0.57 ± 0.06 | 0.33 ± 0.17 | 0.50 ± 0.10 | 0.30 ± 0.18 |
| BASO (%) | Buffer | 0.13 ± 0.03 | 0.13 ± 0.03 | 0.13 ± 0.03 | 0.10 ± 0.00 |
| | mRNA | 0.17 ± 0.03 | 0.10 ± 0.00 | 0.20 ± 0.00 | 0.20 ± 0.00 |
| PT (s) | Buffer | 9.60 ± 0.06 | 9.43 ± 0.03 | 9.40 ± 0.06 | 9.60 ± 0.06 |
| | mRNA | 9.90 ± 0.12 | 9.97 ± 0.06 | 9.83 ± 0.12 | 10.20 ± 0.12 |
| APTT (s) | Buffer | 23.27 ± 0.43 | 23.40 ± 0.35 | 24.03 ± 0.78 | 23.67 ± 0.50 |
| | mRNA | 21.60 ± 1.00 | 21.77 ± 1.06 | 21.80 ± 1.33 | 22.13 ± 1.07 |

Supplementary Table S2. Blood chemistry following repeated intradermal injections in Cynomolgus Monkeys using PYRO

| | Injection | Day -7 | Day 20 | Day 41 | Day 55 |
|---|---|---|---|---|---|
| AST (U/L) | Buffer | 39.33 ± 7.84 | 40.67 ± 11.72 | 31.33 ± 2.91 | 28.67 ± 2.33 |
| | mRNA | 28.00 ± 10.33 | 25.33 ± 13.69 | 24.33 ± 5.46 | 23.33 ± 4.62 |
| ALT (U/L) | Buffer | 57.33 ± 19.41 | 58.33 ± 19.01 | 49.00 ± 17.69 | 56.67 ± 22.93 |
| | mRNA | 45.67 ± 21.79 | 43.67 ± 20.99 | 35.00 ± 17.09 | 34.67 ± 23.79 |
| LD (U/L) | Buffer | 343.7 ± 41.7 | 318.3 ± 32.3 | 294.3 ± 23.3 | 280.3 ± 12.9 |
| | mRNA | 381.0 ± 15.0 | 345.0 ± 17.5 | 317.0 ± 22.4 | 327.0 ± 12.7 |
| CK (U/L) | Buffer | 143.7 ± 11.6 | 157.3 ± 9.0 | 206.7 ± 74.7 | 151.3 ± 26.8 |
| | mRNA | 151.7 ± 16.7 | 156.3 ± 22.1 | 150.3 ± 82.3 | 137.0 ± 15.0 |
| GLU (mg/dL) | Buffer | 104.7 ± 5.6 | 123.3 ± 25.9 | 97.0 ± 13.6 | 71.7 ± 7.4 |
| | mRNA | 97.3 ± 4.3 | 107.7 ± 27.0 | 137.7 ± 13.4 | 95.3 ± 1.2 |
| BIL (mg/dL) | Buffer | 0.12 ± 0.01 | 0.11 ± 0.01 | 0.17 ± 0.02 | 0.12 ± 0.03 |
| | mRNA | 0.11 ± 0.01 | 0.09 ± 0.01 | 0.12 ± 0.02 | 0.09 ± 0.01 |
| UN (mg/dL) | Buffer | 20.10 ± 1.39 | 21.10 ± 1.31 | 18.73 ± 0.90 | 20.20 ± 1.10 |
| | mRNA | 17.93 ± 1.77 | 18.47 ± 2.14 | 16.90 ± 1.76 | 18.60 ± 1.66 |
| CRE (mg/dL) | Buffer | 0.70 ± 0.03 | 0.70 ± 0.08 | 0.67 ± 0.08 | 0.76 ± 0.09 |
| | mRNA | 0.67 ± 0.03 | 0.71 ± 0.04 | 0.65 ± 0.05 | 0.68 ± 0.05 |
| CHO (mg/dL) | Buffer | 118.7 ± 6.1 | 119.0 ± 7.6 | 119.7 ± 6.9 | 109.3 ± 10.7 |
| | mRNA | 131.7 ± 6.4 | 113.3 ± 8.7 | 118.0 ± 7.4 | 111.0 ± 12.5 |
| TG (mg/dL) | Buffer | 44.00 ± 0.58 | 47.00 ± 5.29 | 34.67 ± 6.33 | 45.00 ± 8.96 |
| | mRNA | 23.67 ± 6.03 | 21.00 ± 9.82 | 24.33 ± 6.44 | 24.67 ± 7.62 |
| PL (mg/dL) | Buffer | 200.3 ± 7.2 | 201.0 ± 4.6 | 185.0 ± 6.7 | 189.3 ± 17.8 |
| | mRNA | 185.3 ± 11.0 | 159.7 ± 16.2 | 160.3 ± 10.1 | 166.3 ± 17.8 |
| IP (mg/dL) | Buffer | 4.17 ± 0.31 | 3.55 ± 0.58 | 4.89 ± 0.48 | 3.74 ± 0.40 |
| | mRNA | 4.68 ± 0.25 | 4.97 ± 0.61 | 5.20 ± 0.47 | 4.42 ± 0.18 |
| CA (mg/dL) | Buffer | 9.57 ± 0.25 | 9.34 ± 0.32 | 9.38 ± 0.32 | 9.39 ± 0.34 |
| | mRNA | 9.35 ± 0.34 | 9.24 ± 0.30 | 9.51 ± 0.31 | 9.28 ± 0.24 |
| NA (mEq/L) | Buffer | 149.9 ± 1.0 | 147.4 ± 1.4 | 147.7 ± 1.1 | 149.3 ± 1.9 |
| | mRNA | 149.9 ± 1.2 | 148.8 ± 1.1 | 149.3 ± 1.1 | 149.5 ± 1.5 |
| K (mEq/L) | Buffer | 4.16 ± 0.10 | 3.74 ± 0.15 | 3.88 ± 0.16 | 3.81 ± 0.08 |
| | mRNA | 4.27 ± 0.08 | 4.15 ± 0.18 | 4.13 ± 0.14 | 3.99 ± 0.08 |
| CL (mEq/L) | Buffer | 107.5 ± 0.4 | 105.1 ± 1.3 | 107.8 ± 1.0 | 108.5 ± 1.1 |
| | mRNA | 112.8 ± 1.7 | 109.9 ± 3.1 | 111.8 ± 2.5 | 111.1 ± 1.3 |
| TP (g/dL) | Buffer | 7.33 ± 0.05 | 7.09 ± 0.07 | 7.15 ± 0.04 | 7.09 ± 0.11 |
| | mRNA | 6.93 ± 0.24 | 6.56 ± 0.30 | 6.56 ± 0.21 | 6.61 ± 0.22 |
| ALB (g/dL) | Buffer | 4.17 ± 0.02 | 3.99 ± 0.10 | 4.03 ± 0.09 | 4.06 ± 0.14 |
| | mRNA | 4.06 ± 0.02 | 3.90 ± 0.11 | 3.90 ± 0.06 | 4.03 ± 0.11 |
| A/G | Buffer | 1.32 ± 0.01 | 1.29 ± 0.05 | 1.30 ± 0.07 | 1.34 ± 0.06 |
| | mRNA | 1.43 ± 0.11 | 1.48 ± 0.10 | 1.46 ± 0.06 | 1.57 ± 0.11 |

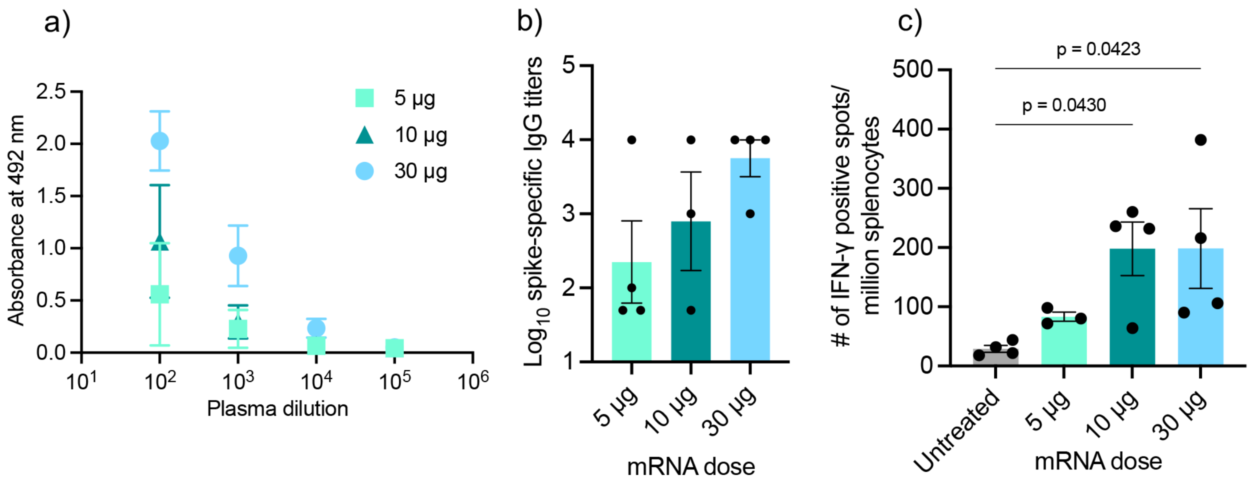


**Supplementary Figure S1.** Humoral and Cellular immunity of naked spike mRNA injected using PYRO in C57BL/6J mice. Mice were injected in a prime-boost setting at a 3-week interval. 2 weeks after the boost, blood plasma and splenocytes were collected for evaluating vaccination effects. (a) Anti-spike IgG ELISA absorbance vs. plasma dilution curves. (b) Log-transformed IgG titers. (c) Quantification of spike-specific IFNγ-positive splenocytes. Data represent the mean ± SEM (n=4). One way ANOVA followed by Dunnett's test.

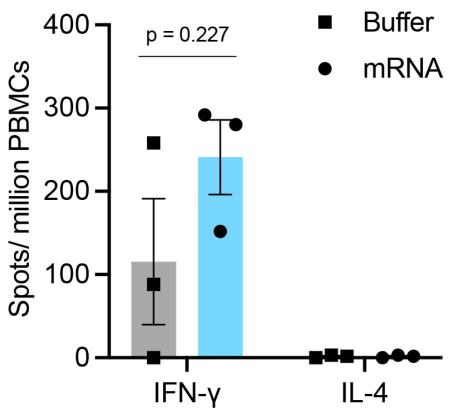


**Supplementary Figure S2.** Cellular immunity of naked spike mRNA injected using PYRO in Cynomolgus Monkeys. ELISpot of PBMCs was performed at day 55. Data represent the mean ± SEM (n=3).
